# Comparative assessment of methods for determining adiposity and a model for obesity index

**DOI:** 10.1101/710970

**Authors:** David Adedia, Adjoa A. Boakye, Daniel Mensah, Sylvester Y. Lokpo, Innocent Afeke, Kwabena O. Duedu

## Abstract

**Background:** Obesity is increasingly becoming a pandemic considering the many risks it pose to other disease conditions. Obesity is largely a measure of adiposity, however, adiposity is not centralized in the human body. This makes it difficult for any single method to adequately represent obesity and by extension the risks specific areas of adipose accumulation pose to specific disease conditions.

**Subjects/Methods:** We evaluated the prevalence of obesity in a cohort of Ghanaian women using the body mass index (BMI) and further sought to evaluate how it compares to other methods of estimating adiposity and the suitability of any particular methods representing obesity in general. We used anthropometry and bioimpedance derived measures of adiposity and derived other indices such as the abdominal volume index (AVI), body adiposity index (BAI) and conicity index (CI).

**Results:** Waist and hip circumference, body fat (%BF) and visceral fat (VF) were positively correlated to BMI whereas skeletal muscle mass was negatively correlated. Physical activity indices did not show any significant correlation with BMI. Prevalence of obesity was 16% and 31% using BMI and %BF respectively. Receiver operating characteristic analysis showed that whereas BMI is effective in predicting underweight, normal weight and obesity it was a poor predictor of overweight.

**Conclusions:** There was also no single measure that could adequately predict obesity as an accumulation fat. Hence, we developed and propose a model as a factor of BAI, %BF, VF and BMI. This model should correctly represent a person’s adiposity status and hence should be evaluated in large cohort studies.

## Introduction

Chronic diseases such as diabetes, hypertension, and metabolic syndrome are rapidly taking over as the major causes of morbidity and mortality in sub-Saharan Africa [1, 2]. The chronic disease burden is attributed to lifestyle changes such as diet, tobacco use and urbanization [2]. In sub-Saharan Africa, the prevalence of infectious diseases such as malaria, HIV, tuberculosis and neglected tropical diseases remains sturdy thereby inflicting a heavy blow on health systems [3, 4]. With the rapidly increasing prevalence of chronic diseases, the health systems will be affected by the rise in infectious diseases co-existing with chronic diseases such as diabetes, hypertension, and metabolic syndrome [5]. Many health systems in the region are under-funded and under-resourced, hence, the chronic disease burden if not nipped in the bud could potentially crash them [6, 7].

Obesity is a widely reported risk factor for chronic diseases such as diabetes, cardiovascular disease and some cancers and recent years have witnessed an alarming increase in the incidence of obesity worldwide [8]. Due to the high health risk associated with obesity, it is important that methods that accurately determines obesity are developed and used. BMI, the ratio of body weight in kilograms to the height in meters squared, has been used to measure obesity for a long while especially in resource limited settings however, BMI measurement does not differentiate between lean and fat mass thus leading to misclassification in some instances. Hence methods that measure direct body fat composition may represent the best standards for determining obesity.

Recent advances in technology have resulted in the development of various tools for measuring adiposity among others. For example, methods like X-ray absorptiometry (DEXA), magnetic resonance imaging (MRI) and bioelectrical impedance analysis (BIA) are available to assess the relative body composition and adiposity. Of these, the BIA methods are relatively cheaper, simple and well adapted for resource-limited settings [9]. The types of BIA instruments have been increasing over time. These instruments can report over 20 parameters on the full body composition including body segment analysis (left arm, right arm, trunk, left leg and right leg), body fat percentage and mass, fat free mass, visceral fat, muscle mass, total body water and body water percentage, among others. However, in many health centres across Ghana lack of the availability of these devices has resulted in the continual use of BMI to predict obesity.

According to the 2016 Global report on diabetes [10], the prevalence of obesity in Ghana were 4.8% and 10.9% (males 4.9% and females 16.8%) respectively. Alarmingly, the prevalence of overweight was 30.8% (males 21.5% and females 39.9%). The primary method for assessing obesity in Ghana is by the BMI method. It has been reported that compared to white Caucasians and other ethnic groups, the South Asian Population have higher amounts of body fats despite having similar or lower anthropometric values [11, 12]. No studies have compared that of Ghanaians in general or among the different ethnic groups in Ghana. As a starting point we sought to (1) compare anthropometry derived adiposity measures and BIA measurements and (2) to determine how accurately different anthropometric measures of obesity can diagnose obesity. Accurate information on fat and other body composition measures will benefit dieticians and other professionals who assist individuals in weight modification programmes.

## Materials and Methods

### Study Design and Population

We conducted a retrospective analysis comparing the prevalence of obesity and associations between adiposity measures among a cross-section of women in Ho, Ghana. The data was collected as part of a community-based Healthy Eating Advocacy Drive (HEAD) outreach conducted between May and December 2016. Data on anthropometric characteristics included Age, Height (m), Weight (kg), Hip Circumference (cm), Waist Circumference (m), Skeletal Muscle (SM, %), Body fat (BF, %), Visceral Fat (VF) and the Resting metabolism rate (RMR). Body Mass Index (BMI) (kg/m^2^), Waist-to-Hip Ratio, Body adiposity index, Abdominal volume index, Visceral adiposity index and Conicity index were derived from the measurements as alternative methods of BMI for determining adiposity. The study was approved by the research ethics committee of the University of Health and Allied Sciences. A standardized questionnaire was used to collect data on the anthropometric measurements and others.

### Physical activity and adiposity measurements

A standard questionnaire was used to obtain information regarding physical activity such as engagement in sports and exercise, work, leisure, sleep and prescribed weight modification or maintenance programmes. Data on behavioural activity related to alcohol consumption, smoking and eating was also collected and characterized.

The Omron body composition monitor (Omron Healthcare Co., Ltd., Kyoto, Japan) was used to measure weight to the nearest 0.1 kg without footwear. There was no adjustment for clothing. Age and gender were inputted into analyser prior to measurements. The VF, SM, BF and RMR were obtained from the BIA. WC and HC were measured using a measuring tape to the nearest 0.1 cm. The WC measurements were taken at the level of the umbilicus with arms folded across the chest whereas the HC measurements were taken at the maximum circumference over the buttocks.

In addition to the BIA measures of adiposity, anthropometric measures of adiposity were calculated using the following standard formulae:

1. Abdominal Volume Index (AVI) - [13]

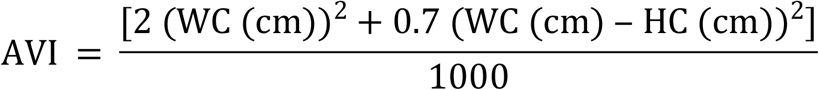
2. Body Adiposity Index (BAI) - [14]

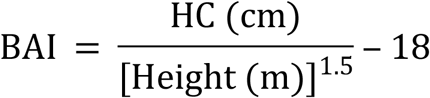
3. Body Mass Index (BMI)

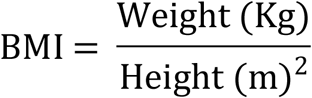
4. Conicity Index (CI) - [15]

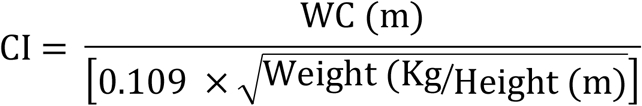
5. Visceral Adiposity Index (VAI) [13]: for females

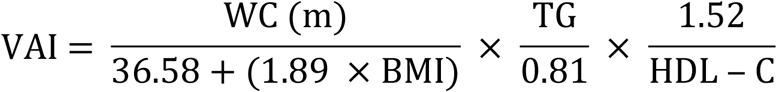

### Statistical analysis

The data was analysed using Minitab version 17 and XLSTAT. We presented the descriptive statistics as (mean ± standard deviation). The strength of linear correlation between BMI and the other alternative method were carried out using Pearson product-moment correlation and Matrix-plot of BMI and other alternative methods. It was very crucial to assess the variation in the alternative methods to BMI for the groups of the BMI status. Analysis of variance concept (ANOVA) was used to test differences between these measures for the four groups of the BMI status. The parametric approach to ANOVA was used for the variables that satisfied both the normality and equal variance assumption, and the variables which did not satisfy these assumptions we applied the non-parametric method (Kruskal-Wallis). The Fisher’s method of multiple comparison was employed for the parametric methods and the Steel-Dwass-Critchlow-Fligner procedure of multiple comparison was employed for the non-parametric approach. All statistical tests were carried out with the level of significance of 5%.

To find other obesity measures, we will use receiver operative characteristics (ROC) with area under the curve (AUC) to assess the classification performance of all candidate obesity measures. For instance, in classifying individuals as obese or not, the probability that higher scores are assigned to people who are obese than people who are not is known as AUC coupled with ROC helps to determine predictive ability of the various measures. When the values of AUC for a particular obesity measure is close to 1, it means that measure can correctly determine if someone is obese or not. The ROC curve is plotted with sensitivity on false positive. Where sensitivity is the probability of a measure classifying someone as obese when the person is actually obese, and false positive is the probability that a person was classified as obese when the person is not obese. These tools have been used by researchers to determine medical tests to discover a particular disease.

We used Multinomial logistic regression to predict probability of underweight, overweight and obesity using body adiposity index, visceral fat and body fat. Confirmatory factor analysis was used to measure the theoretical variable, obesity index.

## Results

We analysed data on 467 women volunteers with a mean age of 46.8±13.3. The prevalence of obesity, overweight and underweight using BMI were 16%, 27% and 9% respectively. Of the data analyzed, 52%, 18%, 15% and 12% were married, single, divorced and widowed respectively with the remainder cohabiting. Twenty percent of the respondents were primary school leavers, 41% were Middle/Junior High School leavers, 10% had secondary school certificate and 12% were tertiary leavers, while the remaining had no formal educational background. Anthropometric indices of the study participants are presented in Table 1.

**Table 1:** Anthropometric characteristics of study participants.

| BMI Category | Number of participants | Age | Height (m) | Weight (kg) | Hip Circumference (m) | Waist Circumference (m) | Skeletal Muscle (%) | Body fat (%) | Visceral Fat Level | Resting metabolic rate |
| --- | --- | --- | --- | --- | --- | --- | --- | --- | --- | --- |
| Underweight | 40 | 49.2±17.9 | 1.59±.100 | 42.8±5.2 | 0.87±0.08 | 0.76±0.10 | 31.9±4.1 | 18.2±7.7 | 2.6±1.4 | 1151±101 |
| Normal | 228 | 45.3±13.4 | 1.58±.059 | 55.1±5.9 | 0.96±0.07 | 0.82±0.10 | 28.7±3.1 | 30.0±5.8 | 5.0±1.4 | 1239±81 |
| Overweight | 124 | 47.8±12.2 | 1.59±.052 | 68.7±5.4 | 1.06±0.06 | 0.94±0.09 | 25.9±2.4 | 39.6±3.8 | 7.9±1.5 | 1376±81 |
| Obese | 75 | 48.3±11.7 | 1.57±.063 | 85.6±14.3 | 1.14±0.09 | 1.01±0.15 | 23.5±3.1 | 46.7±6.3 | 10.5±2.4 | 1511±143 |
| **Total** | **467** | **46.8±13.3** | **1.58±.063** | **62.5±14.7** | **1.01±0.11** | **0.88±0.13** | **27.4±3.8** | **34.2±9.8** | **6.5±2.8** | **1312±146** |

There was a difference in the population mean WC (*p-value<0.0001*) for the BMI categories. Body fat, visceral fat and resting metabolism rate showed significant differences within the BMI categories (*p-value<0.0001*). The relationship between secondary measures of adiposity and BMI are presented in Table 2. WHR for the obese group was significantly higher than all other groups, whilst the mean WHR for the overweight was significantly more than the normal group, it was not significantly different from the underweight group. The population mean AVI and BAI were significantly different for the BMI classifications, and participants with higher values of these measures also had higher BMI. There was no difference in the population mean Visceral adiposity index for the BMI classification. The mean Conicity index for the normal group was significantly lower than the mean Conicity index for the overweight and obese groups. Although the mean conicity index and visceral adiposity index for the underweight group was higher than that of the normal category, there was no significant difference (Table 2).

**Table 2:**
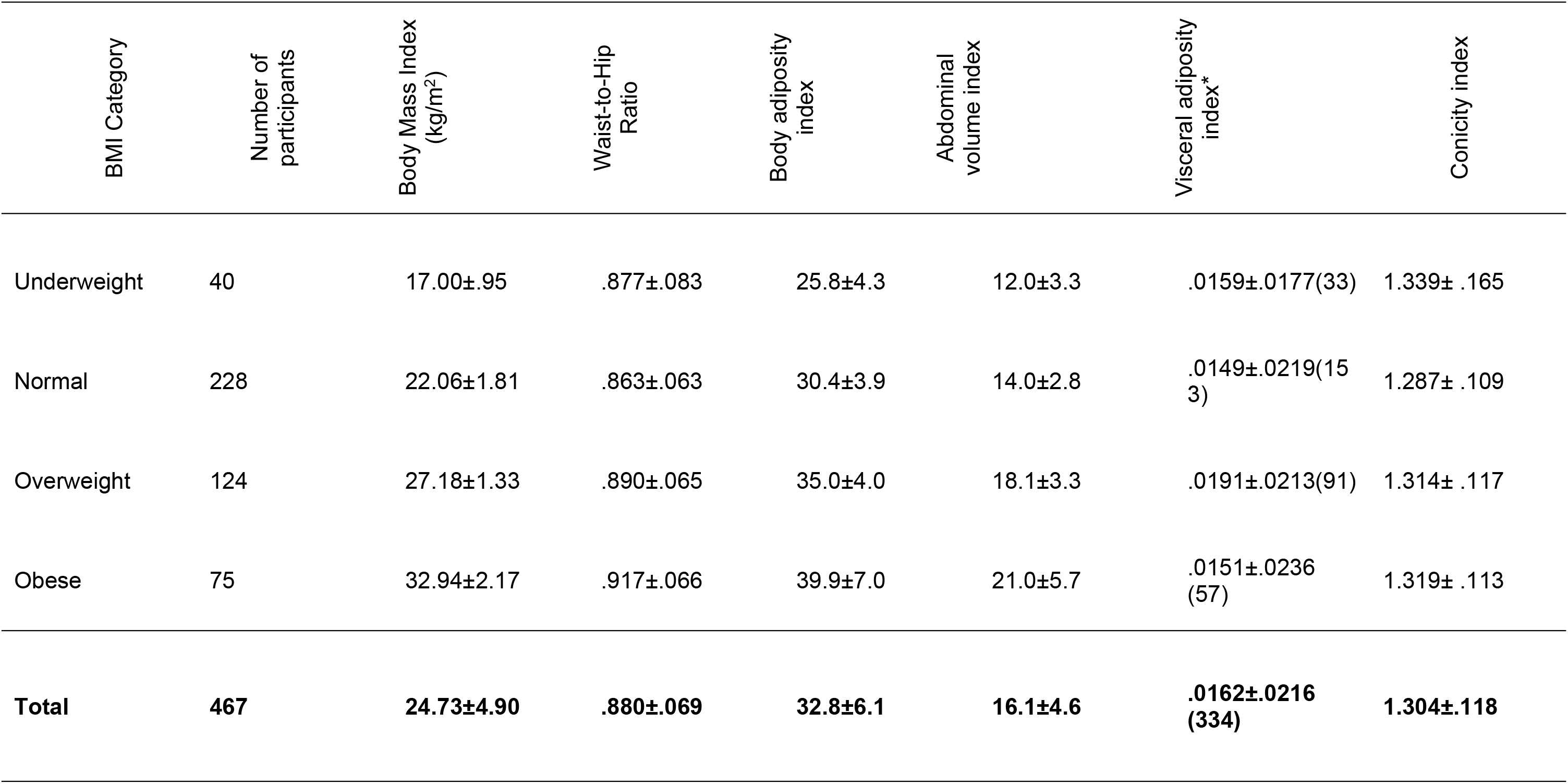
Anthropometry derived adiposity indices of study participants.

| BMI Category | Number of participants | Body Mass Index (kg/m<sup>2</sup>) | Waist-to-Hip Ratio | Body adiposity index | Abdominal volume index | Visceral adiposity index* | Conicity index |
| --- | --- | --- | --- | --- | --- | --- | --- |
| Underweight | 40 | 17.00±.95 | .877±.083 | 25.8±4.3 | 12.0±3.3 | .0159±.0177(33) | 1.339± .165 |
| Normal | 228 | 22.06±1.81 | .863±.063 | 30.4±3.9 | 14.0±2.8 | .0149±.0219(153) | 1.287± .109 |
| Overweight | 124 | 27.18±1.33 | .890±.065 | 35.0±4.0 | 18.1±3.3 | .0191±.0213(91) | 1.314± .117 |
| Obese | 75 | 32.94±2.17 | .917±.066 | 39.9±7.0 | 21.0±5.7 | .0151±.0236(57) | 1.319± .113 |
| **Total** | **467** | **24.73±4.90** | **.880±.069** | **32.8±6.1** | **16.1±4.6** | **.0162±.0216(334)** | **1.304±.118** |

Anthropometric measurements such as body fat, visceral fat, RMR, hip measure, Body Adiposity Index, Abdominal Volume Index, and waist circumference, showed strong positive correlations (R= 0.874, 0.867, 0.804, 0.764, 0.708, 0.667, 0.622 respectively) with body mass index, whilst skeletal muscle had a strong negative linear correlation with BMI (R=−0.685). Waist-to-hip ratio and leisure index showed weak relationships (R=0.283 and −.184 respectively), Visceral adiposity index, conicity index, work index and sports index showed no relationships (R= −0.002, 0.059, 0.043 and 0.062 respectively) with body mass index.

The anthropometric measurements such as body fat, visceral fat, RMR, hip measure, Body Adiposity Index, Abdominal Volume Index, and waist circumference, showed strong positive correlations (R= 0.874, 0.867, 0.804, 0.764, 0.708, 0.667, 0.622 respectively) with BMI, whilst skeletal muscle mass had a strong negative linear correlation with BMI (R=−0.685) (Figure 1). Waist-to-hip ratio and leisure index showed weak relationships (R=0.283 and −0.184 respectively), Visceral adiposity index, conicity index, work index and sports index showed no relationships (R= −0.002, 0.059, 0.043 and 0.062 respectively) with body mass index. Visceral fat followed by body fat had very high correlations with BMI. The correlations within BMI categories showed no relationships for people who were underweight. The category of normal weight had similar correlations as non-classified BMI correlations in Fig 1 whilst the overweight and obese categories showed weak correlations with BMI (Fig 2).

**Figure 1.**
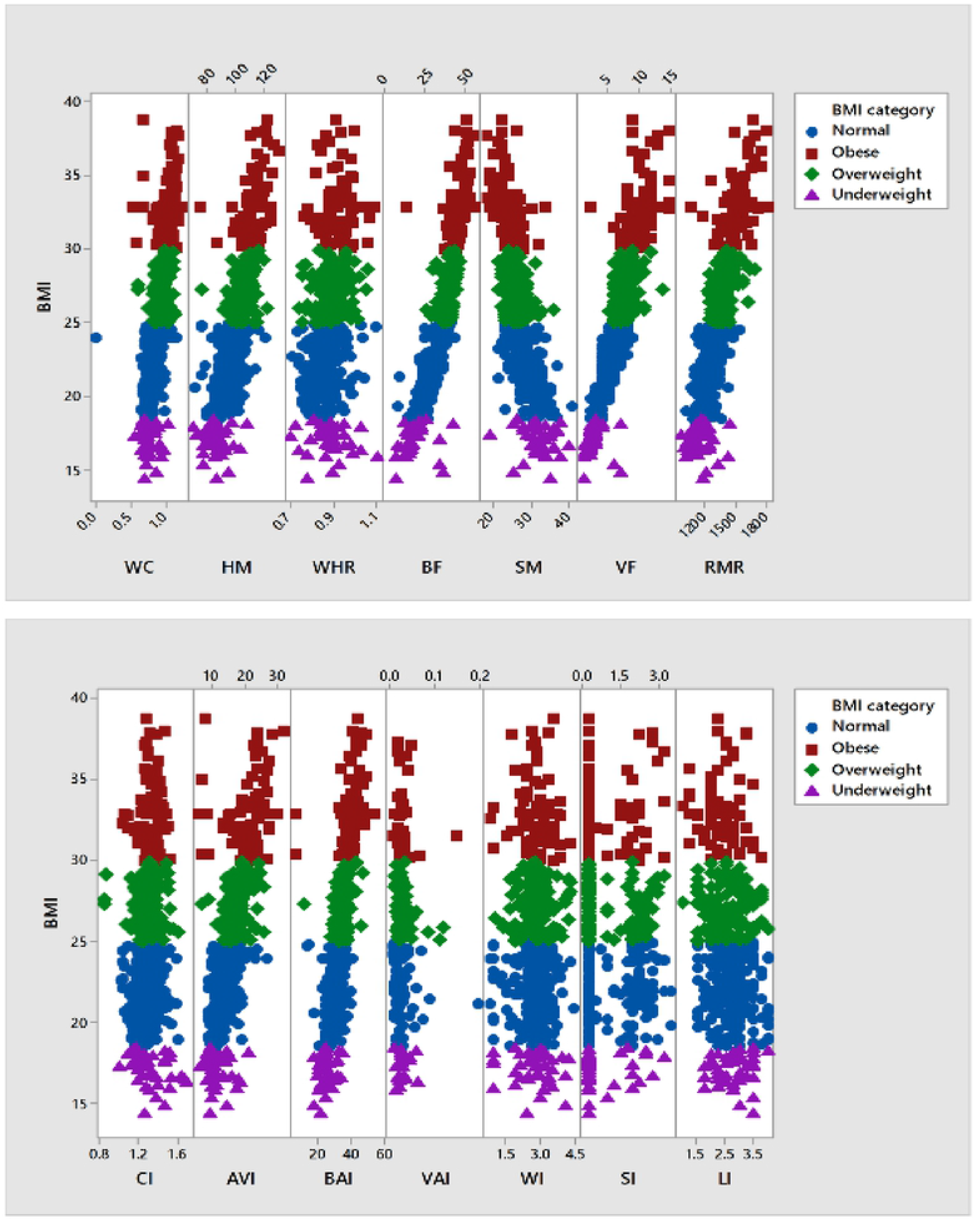
Correlations between BMI and other indices of obesity classified by BMI status. The correlations between BMI and waist circumference (WC), Hip measure (HM), waist to hip ratio (WHR), Body fat (BF), Skeletal Muscle (SM), Visceral fat(VF), Resting metabolic rate (RMR), Conicity Index (CI), Abdominal volume index (AVI), and Body adiposity index (BAI), Visceral adiposity index (VAI), work index (WI), sport index (SI) and leisure index (LI) classified by BMI category is demonstrated.

**Figure 2.**
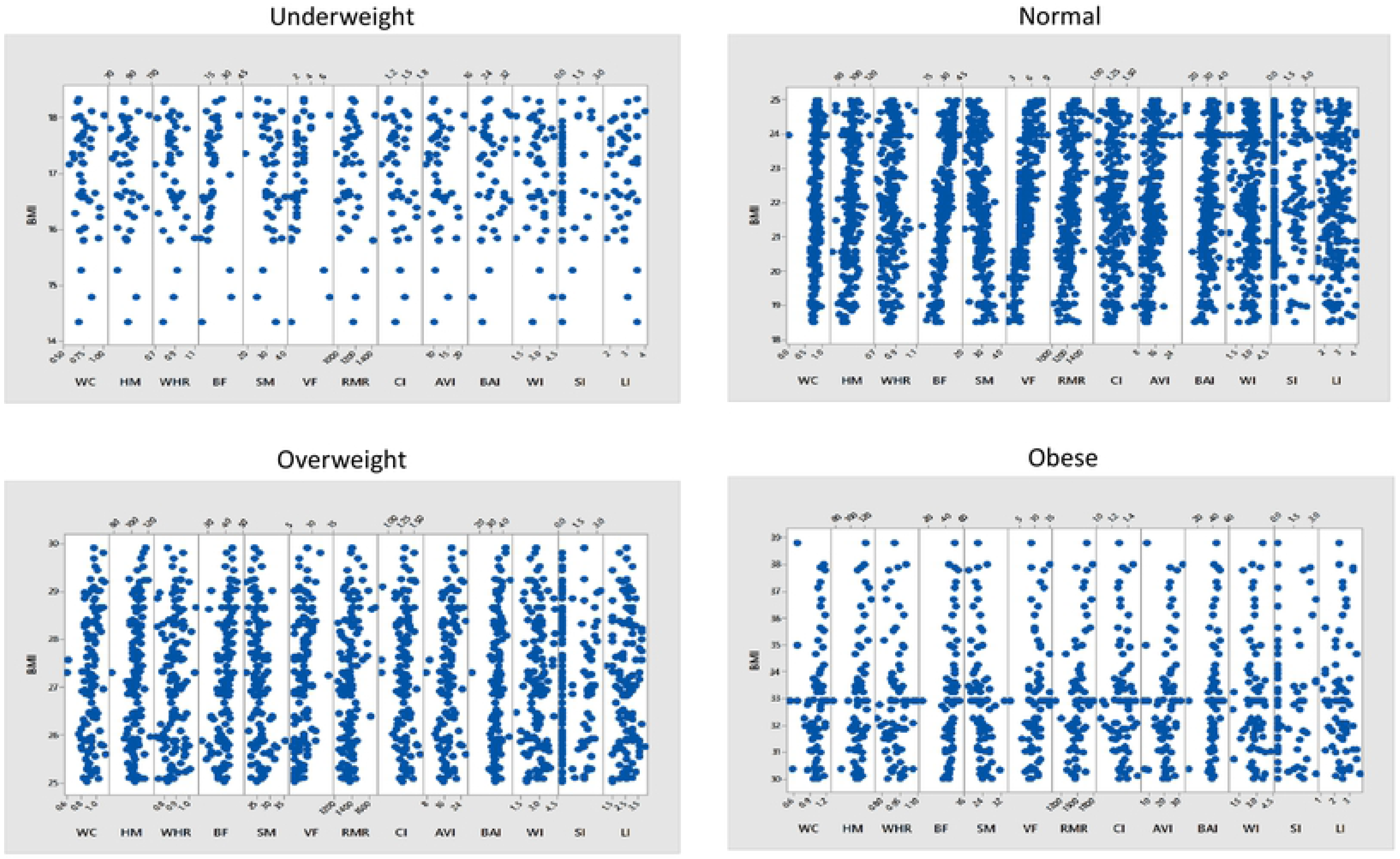
Correlations within different BMI categories and adiposity indices. Figure 2 shows the individual scatter plots (Underweight, Normal, Overweight and Obesity) between BMI and the remaining candidate obesity measures.

To determine the suitability of predicting obesity by the different indices of adiposity using BMI as the gold standard, a receiver operating characteristic analysis was performed. Our results show that measures such as body fat and visceral fat were excellent predictors of adiposity and except for waist to hip ratio all others showed moderate predictive abilities (Fig 3). Table 3 gives a summary of the results in Fig 3. To determine the suitability of predicting obesity by the different indices of adiposity using %BF on the other hand as the gold standard, a receiver operating characteristic analysis was also performed. Measures such as BMI and visceral fat were excellent predictors of low %BF (eg. underweight) and very high %BF (eg. obesity). They were also moderate predictors of normal %BF and a poor predictor of high %BF (overweight). All the other measures of adiposity performed moderately for all the adiposity classes except of overweight.

**Figure 3.**
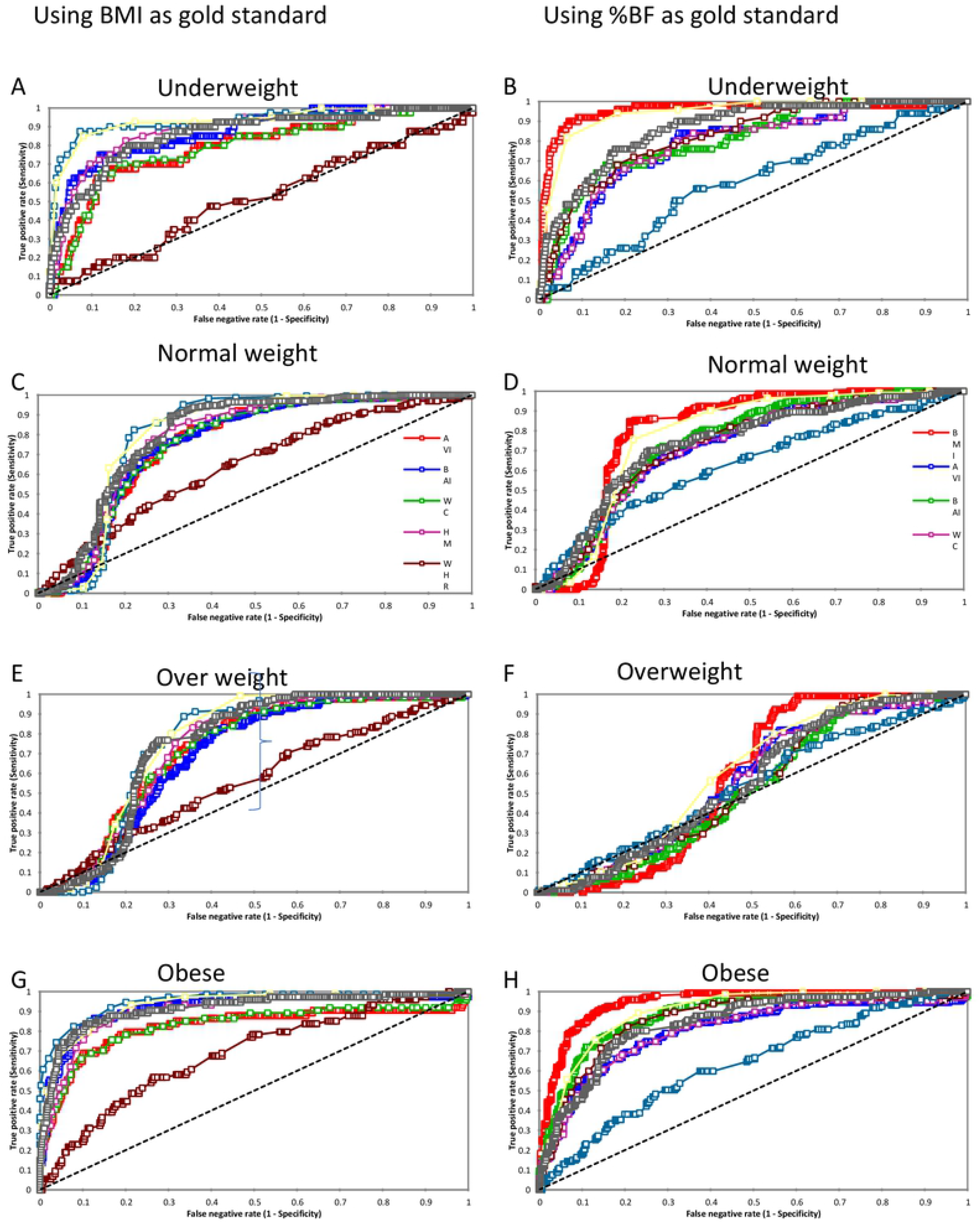

**Figure 4.**
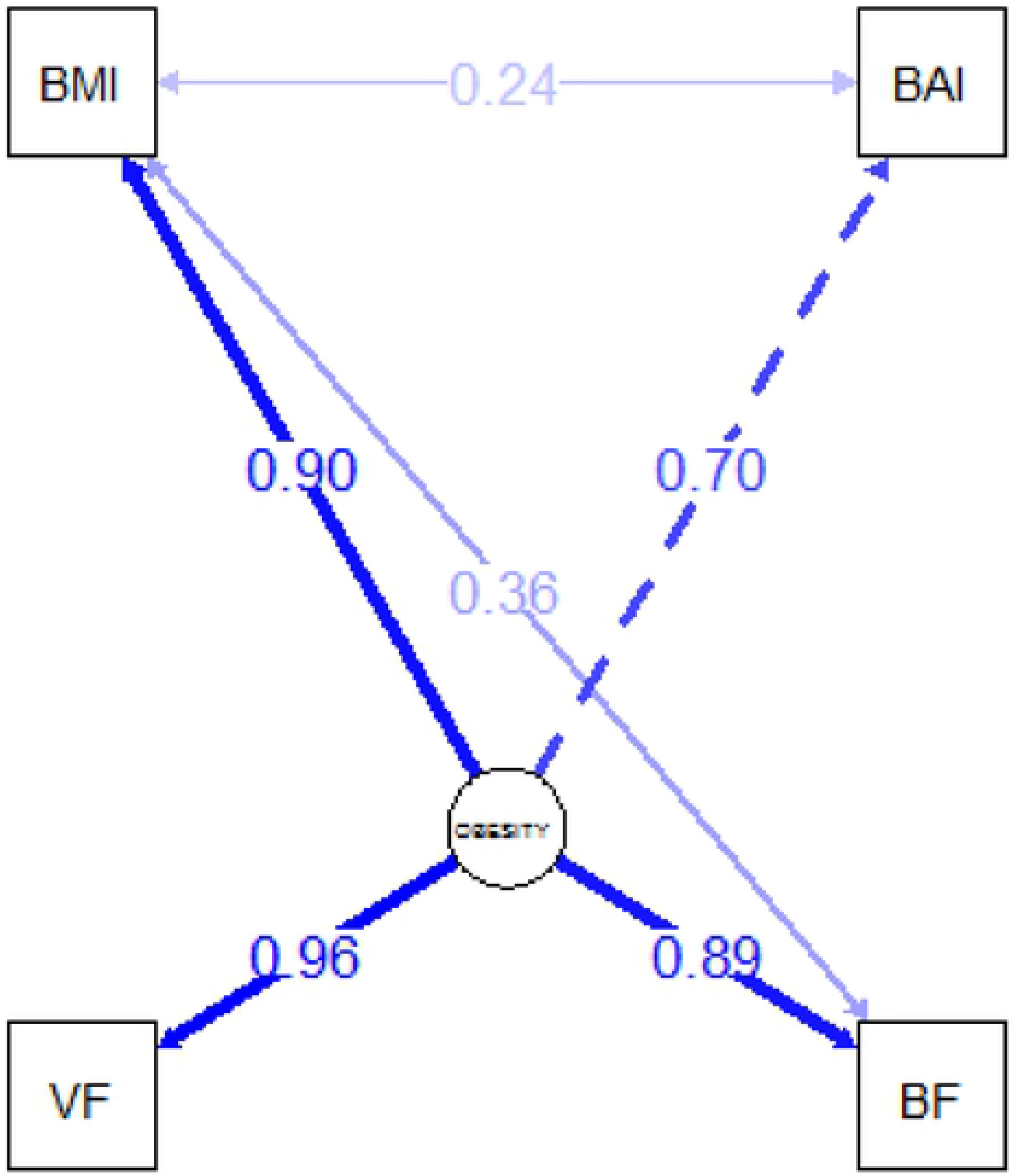

## Discussion

Several large longitudinal studies have shown that obesity is associated with increased risk of chronic diseases such as cardiovascular disease, cancers and diabetes amongst others and that weight reduction reduces the risk of these diseases[16]. In recent years we have witnessed an alarmingly increase prevalence of non-communicable diseases (NCD) [17]. This increased NCD burden is associated with increase in the prevalence of obesity [16]. Thus, reducing the obesity epidemic may in part be an effective tool to solving this increased prevalence of NCD. The gold standard in diagnosing obesity as well as the relationship between the different measures of obesity and other chronic diseases remains debatable. Studies within Mexican and Caucasians population have reported that prevalence of obesity in their study population differ depending on whether the classification was done with BMI or %BF [18, 19]. In a study among Australia women they showed that waist circumference and waist-to-hip ratio were better predictors of cardiovascular disease risk than BMI [20] whereas in a Nigerian population study, BMI in addition to both waist circumference and waist-to-hip ratio were excellent predictors of cardiovascular risk [16]. Lichtash and colleagues tried to find an alternative to BMI as obesity and overweight indicator since BMI which is the most widely used measure does not differentiate between skeletal muscle mass and fat mass. They found that even though BAI had association with some cardiometabolic trait it did not outperform BMI in that regard [21]. These results calls for population based studies that looks at the effectiveness of the different adiposity measure in predicting obesity and risk of obesity-associated

In our study, we assessed how other measures of adiposity compared with traditional BMI and found that some of the measures have strong association with BMI which is an indication that they could be used as well or in place of BMI for assessing obesity. Generally, these measures showed an increasing trend as one progresses from underweight to obesity except for skeletal muscle mass that showed a decreasing trend and this trend was similar to what had been observed previously[16]. Of particular interest is the fact that for measures like WHR, VAI and CI the underweight individuals showed higher levels than normal weight individual even though it did not achieve statistical significance. Although we cannot assign a particular reason to this observation it may be a reflection that although obesity representing an “overfed state” poses a significant health problem undernutrition represented by underweight is also a significant health risk.

The lack of association between physical activity and obesity in our study cohort could be due to the fact that most of these individuals had very similar levels of physical activity hence, sedentary lifestyle may not be the driver of obesity in this population. Further studies looking at dietary habits, genetics and lifestyle choices may be important in driving at the mechanisms underlying obesity in this study population. The results implied that visceral fat followed by body fat had very high correlations with BMI, which means they could be used in place of BMI or possible measures of obesity. The observation suggest that in resource limiting settings BMI may still be effective in predicting obesity however we wanted to test how specific or selective BMI is in making these predictions.

Using BMI as the standard, it was observed that % body fat and visceral fat levels were excellent in predicting underweight, normal weight, overweight and obesity individually. Other measure such as Body Adiposity Index, Abdominal Volume Index, resting metabolic rate and waist circumference reported higher values implying they are able to correctly classify persons within the various BMI classes. The other measures used to predict adiposity showed moderate predictive value except waist to hip ratio (WHR) which showed low predictive value. This results is similar to what has been previously reported among adolescents[22].

Based on these observations we wanted to see how BMI will fare against the gold standard which is %BF. From the AUC values we see that BMI is an excellent predictor of underweight, normal weight and obesity however it was a bad predictor of overweight. This is particularly due to the low specificity even though it has a high sensitivity (Table 3). This observation is problematic since overweight represent pre-obese state and therefore having an accurate measurement is important in preventing the obesity epidemic. Based on the limitation of the sample size for this study we cannot draw conclusion thus large-scale epidemiological studies need to be carry out to ascertain the real association between these parameters. Likewise, the prevalence of obesity increased from 16% to 31% when %BF was used. The results from this study and those from others that have shown presence of masked obesity with the use of BMI, suggests that the current WHO classification for BMI may not be an accurate predicator of obesity in all population and in resource limited settings where BMI is used population specific cut-off points may have to be developed.

Obesity is closely associated with myriad comorbid complications making it increasingly a major socioeconomic burden. Furthermore, specific indices of obesity often suggest risks to specific comorbid conditions. For example, WC and WHR have been described as indices suggesting a higher risk to cardiovascular diseases [20]. Furthermore, visceral fat accumulation has also been described as a risk factor for prostate cancer[23] and metabolic syndrome[24] among others. Hence, there is a need for a central index of obesity that takes into consideration the various indices of adiposity to represent obesity as a health risk. In order to arrive at a model for predicting obesity within our cohort we used model fitting. The fit indices reported showed that the model hypothesized is a good model and can be used to explain the variation in the dataset. The results showed that body adiposity index, body fat, visceral fat and body mass index are the key indices for measuring obesity. The obesity index derived for the population of women in Ho is given by the expression:

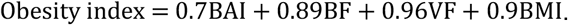

In conclusion our study determined the various adiposity measures that could be used individually or together to assess obesity: Body mass index, body adiposity index, visceral fat and body fat. From the confirmatory factor analysis, visceral fat contributed the highest in the obesity index, followed by body mass index, body fat and body adiposity index. We also observed that although BMI was an excellent predictor of underweight, normal weight and obesity it wasn’t a good predictor of underweight thus we recommend that %BF be used in predicting obesity. We also suggest that ethnic specific cut-off for BMI be developed.

## Acknowledgements

The authors acknowledge all study participants for agreeing to participate in the study. We also acknowledge the Church of Pentecost, Volta Area and the Presbyterian Church of Ghana, Ho District for providing the platform for recruitment of study participants. KOD Acknowledges postdoctoral funding from the African Partnership for Chronic Disease Research (APCDR) towards this work.

## Competing Interests

None declared

## Author Contributions

DA and AAB analysed data and drafted the manuscript. DM, SYL and IA collected and analysed samples and data. DM and SYL contributed to analysis of the data and drafting of the manuscript. KOD conceptualized and designed the study and reviewed the manuscript. All authors approved the final manuscript.

